# Mutant *Zp1* results in Zona Pellucida Lacking and Female Infertility in Rats and Humans

**DOI:** 10.1101/825018

**Authors:** Chao Lv, Hua-Lin Huang, Yan Wang, Tian-Liu Peng, Hang-Jing Tan, Ming-Hua Zeng, Ru-Ping Quan, Hong-Wen Deng, Hong-Mei Xiao

## Abstract

Zona pellucida (ZP) plays a vital role in reproductive processes including oogenesis, fertilization and preimplantation development of embryo. The ZP of humans is composed of four glycoproteins (ZP1-ZP4), same as rats ZP. Our previous research reported a first case of human infertility due to *ZP1* mutation, but the mechanism was unclear. Here we developed a genome editing *in vivo* rat model and a co-transfected *in vitro* cell model to investigate the pathogenic effect. In rat homozygous for the homologous mutation, ZP were absent in all of collected eggs. Further the growing and fully grown oocytes in the mutant ovaries completely lack a ZP but with detectable intracellular ZP1 protein. After mating with male rats, none of the mutant female rats got pregnant. Moreover, the co-transfected cell experiments and the ovarian experiments showed that the truncated ZP1 sequestered intracellularly ZP3 and ZP4 to impede their release outside, resulting in an intracellular accumulation of ZP1, ZP3 and ZP4, leading to absence of ZP in mutant oocytes. Our results clearly establish the causal role of *ZP1* mutation on ZP defects and female infertility.

**Summary statement:** Rat model mirrored completely the phenotypes observed in humans, infertility and abnormal eggs that lack a zona pellucida, through the negative effects of *ZP1* mutation.

## INTRODUCTION

The mammalian zona pellucida (ZP) is a glycoprotein matrix surrounding oocytes, eggs and embryos up to the time of blastocyst hatching (Fig. 1A) (Wassarman, 2008). It plays an important role in production of oocytes, in recognition of gametes, in induction of acrosome reaction, in prevention of polyspermy and in protection of the embryo during its transfer through the fallopian tube (Gupta et al., 2009; Matzuk et al., 2002). When oocytes enter the growth phase, the *ZP* genes were activated early or late, encoding the zona glycoproteins to form the ZP, which is critical for mature, ovulation and fertilization of oocytes *in vivo* (Avella et al., 2014; Wassarman, 1999). During follicular development, the ZP can be first observed as extracellular patches surrounding oocytes in primary follicles which then gradually increase in width around growing oocytes (Gook et al., 2008).

**Fig. 1.**
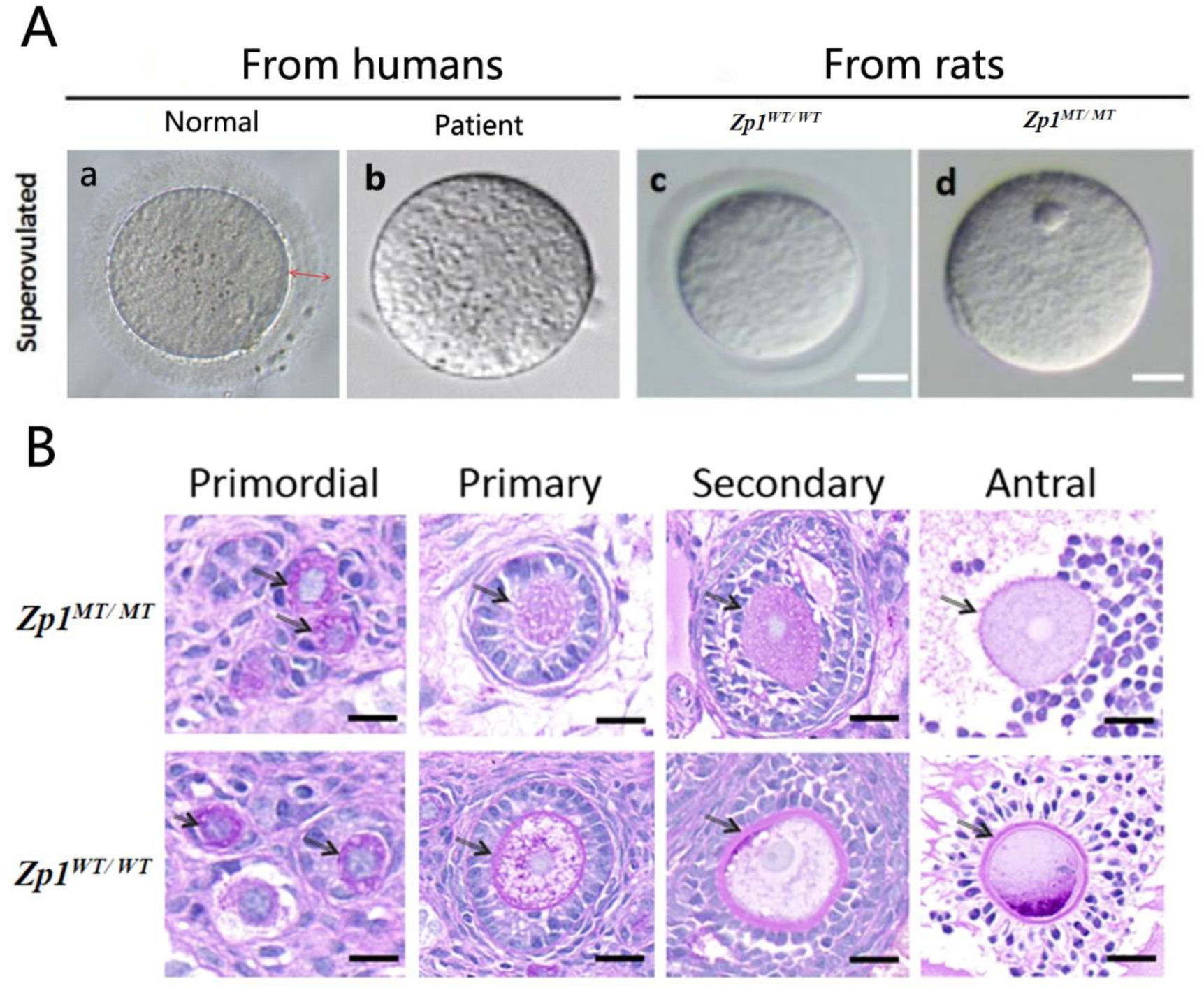
Morphological analysis of ZP in human and rat models. **A**, oocytes from humans, which were cultured and separated from granulosa cells (left). The morphology of the ZP in ovulated eggs from rats under a light microscope: eggs of wild-type rats and mutant eggs lacking a ZP (right). **B**, High magnification images of oocytes at different stages of follicular development from *Zp1*^*MT/MT*^ and *Zp1*^*WT/WT*^ rats. Arrows point to ova. Following the primary follicle stage, the oocytes showed a disorganized follicular structure with the absence of the zona pellucida in mutant oocytes; the antral oocytes were especially clean without any nearby granulosa cells.

Our previous study reported a form of familial infertility with an autosomal recessive mode of inheritance (OMIM 615774), characterized by eggs lacking a ZP (Fig. S1) (Huang et al., 2014). We determined that a frameshift mutation (I390fs404X) in the *ZP1* gene was co-segregated with the ZP defects and infertility in the family. *ZP1* encodes one of the four zona glycoproteins in humans, which is vital for the formation of zona matrix (Ganguly et al., 2010; Nishimura et al., 2019). Based on both humans and rats ZP were constructed of four zona glycoproteins (ZP1-ZP4) (Boja et al., 2005; Lefievre et al., 2004), we developed a rat model that carried a homologous deletion in the *Zp1* gene and a cell model co-transfected with genes encoding the normal and mutant zona proteins (Ma et al., 2017), to investigate the causal effects of this mutation on ZP defects and infertility.

## Materials and Methods

### CRISPR/Cas9-mediated gene editing

By analyzing the homologous genomic sequences of humans and rats, we determined the 8-bp deletion site in the rat *Zp1* genomic DNA sequence (GenBank accession number MK527841), which is located on chromosome 1. Using VectorBuilder software (www.vectorbuilder.com), we designed a *Zp1* guide RNA (gRNA) and a donor oligo (Table S1) bearing the locus-specific homologous sequence and the intended 8-bp deletion according to the quality scores and off-target analysis (Fig. S2). The Cas9 mRNA, gRNAs generated by transcription *in vitro* and donor oligo were coinjected into fertilized eggs to generate the 8-bp deletion in the rat *Zp1* gene (Table S1). The injected zygotes were transplanted into the womb of foster mothers (Sprague Dawley rat, SD rat) (Sander and Joung, 2014). The founder mice (F0) were obtained in approximately 3 weeks. All animal experiments were approved by Department of Zoology, Central South University. The positive F0 offspring were selected to mate with wild type (WT) rats to generate hybrid F1 offspring, some of which could carry heterozygous *Zp1* mutation with stable inheritance. The F1 heterozygous rats were cross-hybridized with each other to generate additional F2 homozygous mutant type (MT) rats. The *Zp1*^*WT/WT*^ and *Zp1*^*MT/MT*^ female rats were hybridized with *Zp1*^*WT/WT*^ male rats to study their fertility.

### DNA, RNA and protein analysis

The test primers were designed to match the segment of gDNA and transferred anti-cDNA containing the 8-bp deletion (Table S2). Genomic DNAs were obtained from tail biopsy specimens and detected by Sanger sequencing. According to the results of the off-target analysis, the designed primers were located on the promoter region of the *Zp1* gene and the potential off-target sites (Table S3). RNAs from ovarian tissues of the rats were dealt with reverse transcription-polymerase chain reaction (RT-PCR) to subject to polyacrylamide gel electrophoresis (PAGE) analysis and Sanger sequencing analysis to identify the targeted 8-bp deletion.

Glycoproteins from ovarian tissues were extracted from 8-week-old female rats. The ovarian tissues were homogenized with M-PER Mammalian Protein Extraction Reagent (Pierce Biotechnology, Rockford, IL, USA) supplemented with Halt TM Protease Inhibitor Cocktail (Thermo Fisher Scientific). Following centrifugation (16,000 g, 4°C), ovarian lysates were separated by sodium dodecyl sulfate polyacrylamide gel electrophoresis (SDS-PAGE) and transferred onto polyvinylidene difluoride (PVDF) membranes, which were probed with antibodies directed against a peptide that had been mapped near the N-terminus of ZP1 (G-20, Santa Cruz, CA, USA, Fig. S4) or against GAPDH (Santa Cruz) and detected by secondary antibodies (Santa Cruz) with an ECL western blotting kit (Pierce Biotechnology).

### *In Vitro* Studies of ZP

The oocytes collected from the patient in the family were observed under a light microscope (Fig. S1) (Huang et al., 2014). All collection and experiments involving samples from patients with infertility were approved the Institute of Reproduction and Stem Cell Engineering, Central South University. All patients provided written informed consent.

Some 8-week-old female rats were sacrificed 21 hours after injection of human chorionic gonadotropin (hCG, 45 IU), and others were sacrificed 69 hours after injection of pregnant mare serum gonadotropin (PMSG, 45 IU). Oocyte cumulus complexes (OCCs) were recovered from *Zp1*^*MT/MT*^ female rats by ovarian puncture. The OCCs were immersed in hyaluronidase droplets, and the cumulus granule cells were gently removed by pipetting to isolate eggs for observation under a visible light microscope.

To highlight the ZP, periodic acid Schiff (PAS) staining was conducted. Ovaries from 8-week-old females (during the sexual maturity period) expressing one of three genotypes were of similar size and morphological appearance. After fixation in 4% paraformaldehyde solution (16-20 hours, room temperature, RT), sections of paraffin-ovarian tissue chips were sliced to a thickness of 4 μm at 100 μm intervals. The mounted sections were stained with PAS reagent and hematoxylin and scanned with a slice scanner (Pannoramic MIDI, 3D HISTECH).

### Protein-Protein Interaction Analysis

The expression vectors for ZP1^WT^, ZP1^MT^ (I390fs404X), ZP2, ZP3 and ZP4 were cloned and inserted into the eukaryotic expression vector plasmid pENTER. In addition, different tags were fused to the C-terminus of genes into *the* vector plasmid pcDNA3.1 (+) to encode ZP1^MT^-FLAG, ZP1^WT^-FLAG, ZP2-MYC, ZP3-V5 and ZP4-HA to avoid the interference of endogenous proteins from tool cells (Fig. S3). The information for the ZP cDNA sequence was acquired from NCBI (www.ncbi.nlm.nih.gov).

Human embryonic kidney 293T (HEK293T) cells (China Center for Type Culture Collection, Wuhan, China) were grown (37°C, 5% CO2) to 70-80% confluence in dulbecco’s modified eagle medium (DMEM) supplemented with 10% fetal bovine serum (Gibco, Thermo Fisher Scientific, Waltham, MA, USA). Transient transfections were performed with Lipofectamine 3000 (Thermo Fisher Scientific). For each transfection, 6 µl of the Lipofectamine 3000 transfection reagent was added to 125 µl of Opti-MEM (Gibco) to which 2.5 µg of template plasmid had been added (cotransfection of ZP1^MT^ or ZP1^WT^ with ZP2, ZP3 or ZP4) and incubated (5 minutes, RT). The mixture was added to the wells with growing cells. Transiently transfected cells were harvested at 24∼48 hours for analysis.

For coimmunoprecipitation (co-IP) analysis, the proteins from cotransfected cell lysates after 24 hours were precipitated with an antibody against the N-terminus of ZP1 (G-20, Santa Cruz). Then, the collected precipitates were analyzed by immunoblotting (IB) with antibodies raised against ZP2 (C-7, Santa Cruz), ZP3 (H-300, Santa Cruz) or ZP4 (I-14, Santa Cruz) to detect coprecipitated zona glycoproteins (Fig. S4).

For IB analysis of ZP1, ZP2, ZP3 and ZP4, zona proteins with specific tags 48 hours after transfection from the culture medium were enriched by IP with a FLAG-tag antibody for labeling ZP1 (AT0022, CAMTAG, WI, USA), a MYC-tag antibody for labeling ZP2 (AT0045, CAMTAG), an V5-tag antibody for labeling ZP3 (AT0497, CAMTAG) and a HA-tag antibody for labeling ZP4 (AT0046, CAMTAG).

### Fluorescent staining

For immunofluorescence (IF) staining, ovarian tissues were isolated and fixed in 3% paraformaldehyde (3-5 hours or overnight, RT), rinsed and transferred to 70% ethanol. The tissues were dehydrated and embedded in methacrylate and were cut into 4 mm sections, which were rehydrated and rinsed three times with phosphate-buffered saline (PBS). The sections were blocked with 5% bovine serum albumin (BSA) for 1 hour and incubated (1 hour at 37°C) with antibodies directed against ZP1 (D-4, Santa-Cruz), ZP2 (PA5-75949, Abcam, Cambridge, United Kingdom), ZP3 (G-1, Santa-Cruz Science) or ZP4 (PA5-37086, Abcam), respectively, and washed three times with PBS, incubated with Alexa-Fluor 555-conjugated donkey anti-rabbit secondary antibodies (Life Technologies, Ghent, Belgium), washed three times in PBS, and counterstained with DAPI (Technologies). Confocal images were obtained with a microscope (ZEISS LSM 880 + Airyscan, Jena, Germany).

Following the above protocol, cells that were double-transfected with plasmids carrying ZP1^WT^ or ZP1^MT^ and ZP2, ZP3 or ZP4, were examined by confocal laser-scanning microscopy, using Alexa-Fluor 488- and Alexa-Fluor 594-conjugated secondary antibodies (Abcam, Cambridge, United Kingdom).

### Statistical Analysis

For each group, the mean ± standard error (s.e.) was calculated. Kruskal-Wallis Test and chi-square test were used for statistical analysis of enumeration data; significance was assumed at *P*<0.05.

## Results and Discussion

### Targeted mutation in *Zp1* gene

Since ZP of both humans and rats are composed of four zona glycoproteins, while the mice ZP consists of three glycoproteins(ZP1-ZP3), the basic structure of ZP in humans and rats may be more similar than that in mice (Boja et al., 2005; Familiari et al., 2006; Lefievre et al., 2004). Although most previous studies exploring the *in vivo* role of the ZP in fertility have been based on mouse models, particularly those utilizing knockouts of ZP genes(Liu et al., 1996; Rankin et al., 1999; Rankin et al., 2001), we chose rat as the animal model for our study to explore the role of truncated ZP1 protein contributing to infertility.

Based on bioinformatics analysis, wild type ZP1 of rat and humans resemble one another, with 67.9% sequence identity (https://www.uniprot.org/blast/uniprot), and the 8-bp deletion at nucleotides 1174-1181 (TCTTCTCA) of the CDS (Coding sequence) of rat *Zp1* (NM_053509.1) resulted in a premature stop codon expected to give rise to a protein truncated at amino acid 401 (I379fs401X) (Fig. 2A). The genotypes of the mutant rats were confirmed using Sanger sequencing (Fig. 2B). The results of PAGE and sequence also showed the mRNA of *Zp1*^*MT/MT*^ rat carrying the 8-bp deletion (Fig. 2C).

**Fig. 2.**
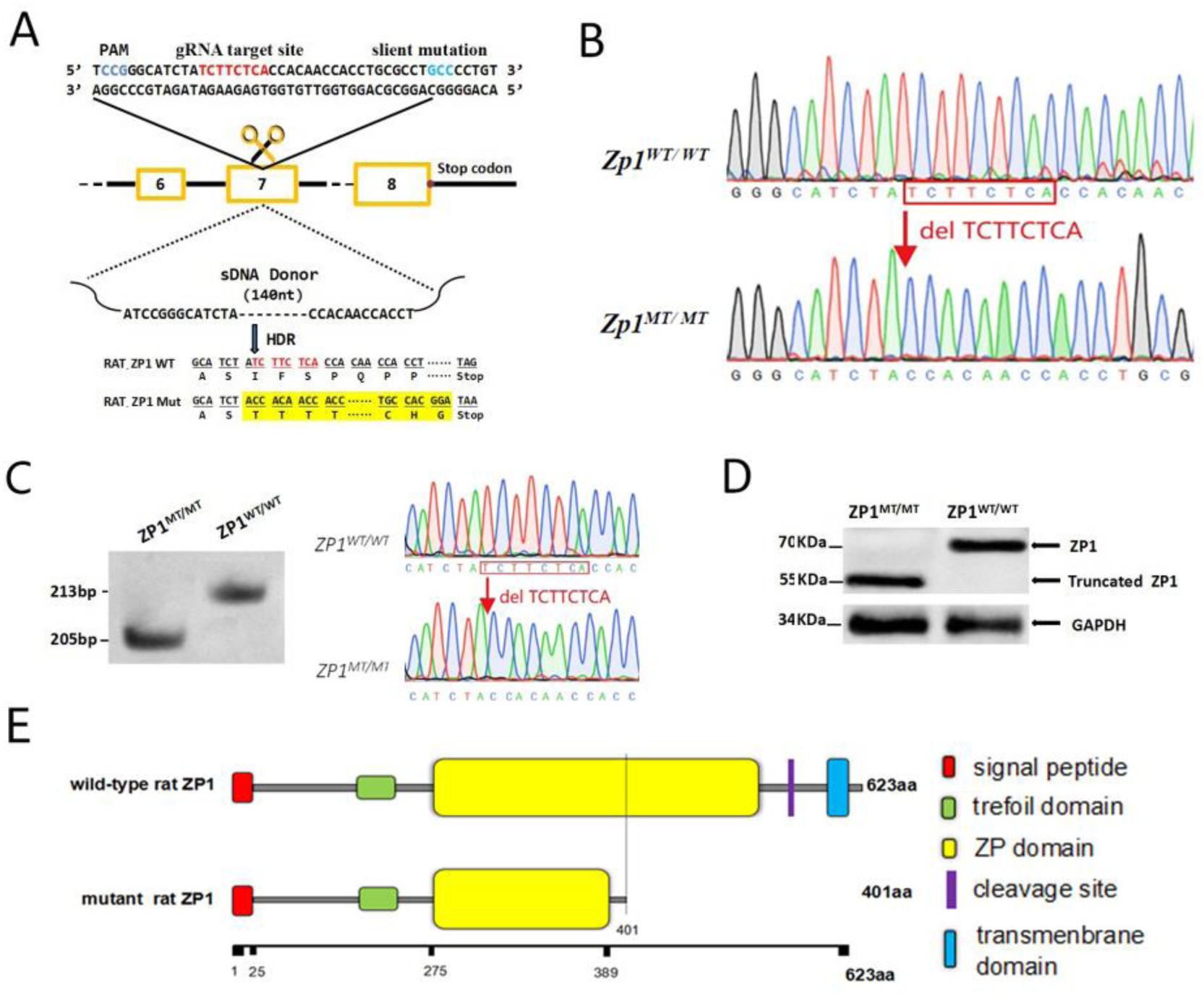
Analysis of *Zp1* mutation in rat models. **A**, the schematic diagram for the generation of mutant Zp1 rats by the CRISPR/Cas9 system. The mutant ZP1 rats showed a homozygous 8bp-deletion at nucleotides 1174-1181 (TCTTCTCA) of rat ZP1 CDS (CCDS37918), leading to a frameshift and the formation of a premature stop codon, I379fs401X. **B**, Genotyping of *Zp1*^*MT/MT*^ rats was performed by PCR and Sanger sequencing of the genomic DNA. Mutant *Zp1* rats exhibited a homozygous mutation [an 8 bp deletion at nucleotides 1174-1181 (TCTTCTCA) of the CDS of rat *Zp1* gene], while a heterozygous mutation was observed in heterozygotes. **C**, The results of PAGE and sequence of Zp1 cDNA in *Zp1*^*MT/MT*^ and *Zp1*^*WT/WT*^ rats. **D**, The ZP1 proteins from the ovaries of *Zp1*^*MT/MT*^ and *Zp1*^*WT/WT*^ rats detected with a special antibody raised against ZP1. GAPDH was used as an internal standard. **E**, The domain organization of wild-type (upper) and mutant (lower) ZP1. The predicted structure shows the deleted portions (e.g., part of the ZP domain in yellow, consensus furin cleavage site in violet, the transmembrane domain in blue and other C-terminal domains).

The results of western blotting to examine the proteins from the *Zp1*^*MT/MT*^ female rats showed a truncated ZP1 protein (Fig.2D), suggesting the retention of the N-terminal domains (e.g., N-terminal signal sequence, the trefoil domain, and the first half of the ZP domain) and the absence of the C-terminal domains (e.g., external hydrophobic patch, consensus furin cleavage site, transmembrane domain, cytoplasmic tail, and the second half of the ZP domain) (Fig.2E) (Ganguly et al., 2010; Gupta, 2018; Monne et al., 2008; Zhao et al., 2003). Similarly, the truncated ZP1 protein of rat, generated for this study by an 8bp deletion, closely resembles the truncated mutant ZP1 that we identified in humans, with 65% sequence identity.

### The homozygous mutant rats were infertile and produced eggs lacking ZP

The fertility of the mutant rats was assessed. Wild-type males were caged with *Zp1*^*WT/W*^ and *Zp1*^*MT/MT*^ females over at least one pregnant cycle in a breeding period of 6 months. Live births occurred 3 weeks later. While *Zp1*^*WT/WT*^ females repeatedly gave birth to normal litter sizes, *Zp1*^*MT/MT*^ female rats never gave birth when caged with several fertile males (p=0.003, Table 1). A continuous breeding period of 12 months was observed for *Zp1*^*MT/MT*^ females, indicating that *Zp1*^*MT/MT*^ females were sterile.

**Table 1.** Fertility of *Zp1*^*WT/WT*^ and *Zp1*^*MT/MT*^ female rats.

| | $Zp1^{WT/WT}$ | $Zp1^{MT/MT}$ |
| --- | --- | --- |
| Pregnancy cycle* | 2.75±0.43 (4) | 0 (6) |
| Litter size* | 11±1.87 (4) | 0 (6) |
| *Average ± s.e. (numbers of animals). |  |  |

In order to get insight into the mechanisms underlying the infertility elicited by ZP1 mutation, oocytes were recovered from oviducts of the two genotypes of female rats after hyperstimulation. In agreement with the infertility of *Zp1*^*MT/MT*^ females, all of the mutant eggs completely lacked a ZP (Fig. 1A). These findings suggest that the significantly infertility from *Zp1*^*MT/MT*^ female rats may be attributed to the eggs without a ZP lost their capacity for *in vivo* fertilization or early embryonic development.

To elucidate the defect in oogenesis of *Zp1*^*MT/MT*^ rats, we performed detailed ovarian morphological analyses. The ovarian sections detected by morphological analysis with PAS staining showed the absence of ZP in mutant homozygous oocytes and disorganized follicular structure at all of the follicular development stages, especially among the antral follicles (Fig. 1B).

The rat model mirrored completely the phenotype observed in our patients (Fig. 1A) (Huang et al., 2014), which is similar to the phenotypes in female mice that lack either *Zp2* or *Zp3 (Liu et al., 1996; Rankin et al., 2001)*, and is more severe than those in mice carrying mutations in *Zp1* gene (Rankin et al., 1999; Wang et al., 2019), suggesting rat may be a more suitable animal model for studying human infertility due to ZP defect.

### The intracellular abnormality of zona proteins leads to ZP lacking

To examine the location and the trafficking of zona pellucida proteins in oocytes, ovarian sections were stained with monoclonal antibodies specific to ZP1, ZP2, ZP3, or ZP4, respectively, and the IF stained specimens were detected with confocal laser scanning microscopy. Mutant ZP1, normal ZP3 and ZP4 were not detected in the extracellular ZP area but were found throughout the cytoplasm in the oocytes of mutant homozygotes, whereas ZP2 was observed only on the cytoplasmic membrane. In contrast, all four zona proteins were detected within the zona matrix of normal growing oocytes but not inside ova (Fig. 3A). As reported earlier, the absence of ZP3 appeared to preclude the formation of a visible extracellular zona matrix. These results suggested that mutant ZP1 cannot be transported out by itself and also affects the transport of ZP3 and ZP4, so their non-release out of ova can cause absence of ZP.

**Fig. 3.**
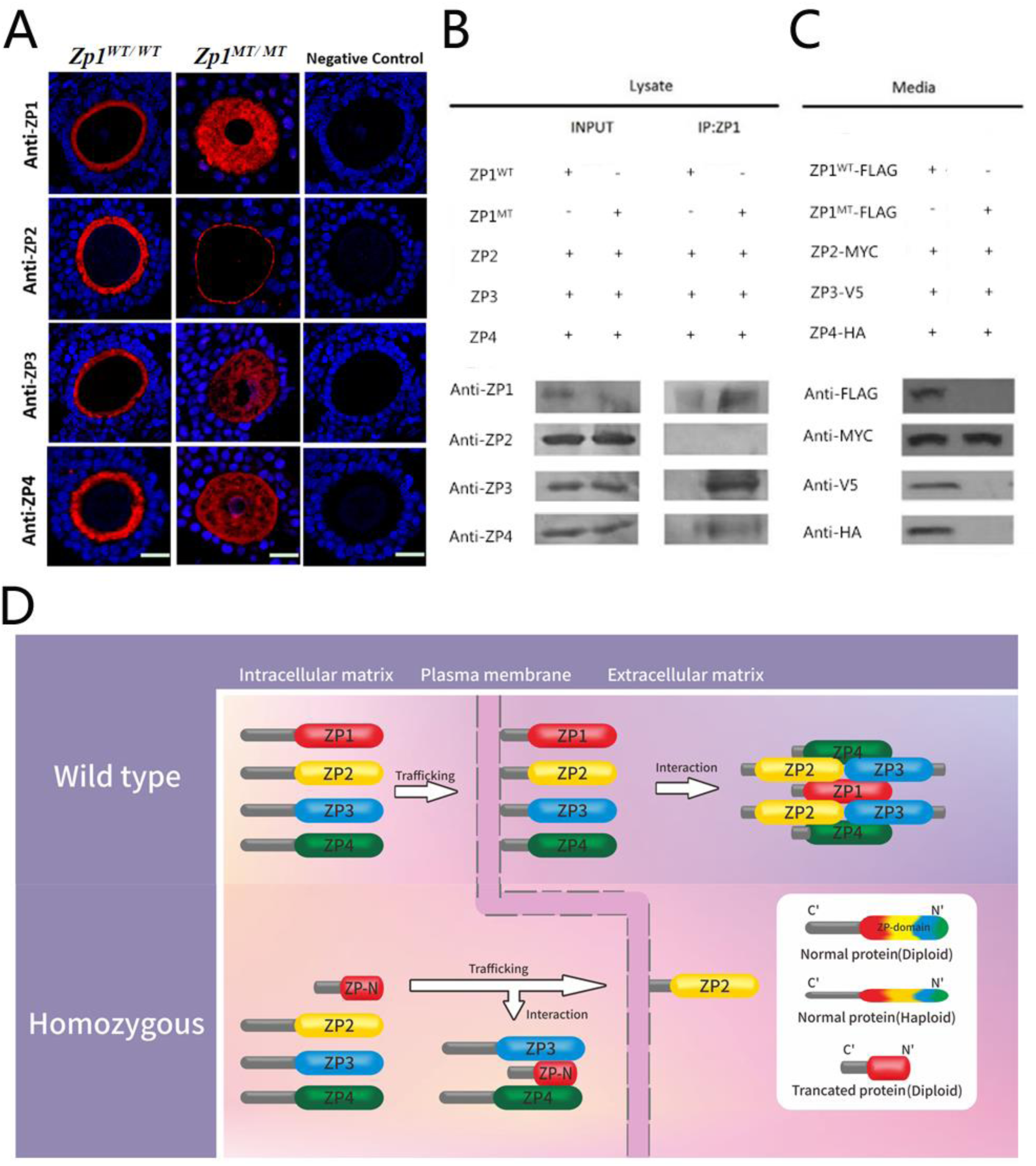
Functional study and Pathogenic model of *Zp1* mutation. **A**, The growing oocytes imaged with the use of confocal microscopy. The ovaries were isolated from *Zp1*^*MT/MT*^ and *Zp1*^*WT/WT*^ rats, fixed, sectioned and stained with antibodies against ZP1 (first row), ZP2 (second row), ZP3 (third row) or ZP4 (fourth row). Fluorescent signals indicating mutant ZP1 were not detected in the extracellular ZP of *Zp1*^*MT/MT*^ rat ovaries but were detected inside the oocytes, which was similar to ZP3 and ZP4, whereas only ZP2 was detected on the cytoplasmic membrane. In normal oocytes, ZP1, ZP2, ZP3 and ZP4 were present in the ZP surrounding the oocytes but were not detected internally. **B**, The interactions of ZP1^WT^ and ZP1^MT^ with ZP glycoproteins. Lanes 1 and 3 show ZP1^WT^ cotransfected with ZP2, ZP3 and ZP4; lane 3 shows IP with an anti-ZP1 antibody; and lane 1 shows the input. Lanes 2 and 4 show ZP1^MT^ cotransfected with ZP2, ZP3 and ZP4; lane 4 shows co-IP with an anti-ZP1 antibody; and lane 2 shows the input. **C**, The media assayed by IP and IB, using monoclonal tagged antibodies against FLAG, MYC, V5 and HA. **D**, The model of the mechanism that prevents the formation of the ZP in homozygotes. ZP1, ZP2, ZP3 and ZP4 are vital normal zona proteins expressed during the formation of a normal ZP in humans. Blue shapes denote wild-type alleles and yellow shapes refer to mutant alleles.

To investigate the functional mechanisms of mutant ZP1 on the intracellular trafficking of zona proteins, co-IP and IF analysis was performed to explore the interactions between ZP1 and other zona proteins. Truncated ZP1 interacted with ZP3 and ZP4 but not ZP2 inside the co-transfected cells, where normal ZP1 did not interact with ZP2, ZP3 or ZP4 (Figs. 3B and S5).

To test the secretion of zona proteins, IB of IP-enriched secreted proteins with specific tags from the medium showed that only ZP2 was detected in the medium from the ZP1^MT^ cotransfected system, while all four ZP proteins were detected in the medium from the ZP1^WT^ cotransfected system (Fig. 3C).

These data indicated that the mutant ZP1 protein prevented the secretion of ZP3, ZP4 and itself out of the ova to interact with ZP2 to form the basal filaments of the zona matrix, resulting in lack of formation of ZP of oocytes (Fig. 3D). These results could be explained by the retention of the partial ZP domain and the absence of cytoplasmic-tail and transmembrane domains in the mutant ZP1 protein that promotes intracellular sequestration of ZP3 and ZP4 proteins, thus impeding their section outside of ova to form the ZP (Jimenez-Movilla and Dean, 2011). The pathogenic mechanism of the familial infertility was identified as a gain of function in recessive mutations that the abnormal protein in homozygotes causes disease by affecting the function of other proteins in addition to the loss of its own function (Hastings et al., 2009; Wilkie, 1994).

*Zp1* mutation can lead to congenital deficiency that loss of ZP which leads to infertility, which is verified in the rat model. The interaction between truncated ZP1 and ZP3 or ZP4 is gained in the cell model, which affects their normal transport and section. Our results suggest that normal ZP1 is crucial for structure of ZP and fertility of oocytes.

## Acknowledgements

This study was supported by the National Natural Science Foundation of China (81471453 and 81501248), the National Key Research and Development Program of China (2016YFC1201805 and SQ2017YFSF080009), and the Natural Science Foundation of Hunan Province of China (2015JJ2166 and 2017JJ3425). We thank Kai Yuan, Fang Chen from the Institute of Molecular Precision Medicine, Xiangya Hospital and the members of the animal center of Central South University.

